# Evolution of Reproductive Plasticity in a Seasonal Tropical Environment

**DOI:** 10.64898/2026.01.20.700078

**Authors:** Marcus Hicks, Zunilda Escalante, Lizett Retuerto, Jamal Kabir, Sridhar Halali, Geoffrey Gallice, Vicencio Oostra

## Abstract

Seasonality drives the evolution of reproductive plasticity in butterflies. An extreme form is reproductive diapause, where individuals halt reproduction during unfavourable seasons through cascades translating predictive environmental cues into reproductive phenotypes. How diapause evolves from milder forms of plasticity through changes in those cascades is poorly understood. In the tropics seasonality is common, but it is unclear how phylogenetically widespread reproductive diapause is, and exactly how seasonal tropical climates drive reproductive plasticity. Here, we study reproductive plasticity in Amazon butterflies using a multi-year monthly time series of reproductive phenotypes. Sampling 4 subfamilies across Nymphalids, we observe a phylogenetically widespread occurrence of diapause, suggesting repeated evolutionary changes in reproductive plasticity. Detailed analyses of two *Catonephele* species reveal a shared temperature response, but dry season diapause only in *C. acontius*. Thus, evolution of diapause combines cue conservation with species-specific divergence of trait plasticity, suggesting gradual and modular evolution of reproductive plasticity.

## 1 | Introduction

Many organisms experience seasonality and have evolved key adaptations to it. In Lepidoptera (butterflies and moths), two main responses to cyclical environmental variability are migration, a spatial response, (Merlin & Liedvogel, 2019; Pollard et al., 1998; Sparks et al., 2005) and diapause, a temporal response (Hoffmann & Bridle, 2022; Merlin & Liedvogel, 2019; Musolin & Saulich, 2012). An important form of diapause is adult reproductive diapause which is particularly important in the tropics (Denlinger, 2022; Denlinger et al., 2012; Halali et al., 2024; Koštál, 2006; Nylin, 2013). This is the halting of oogenesis in females, which allows them to effectively ‘wait out’ poor environmental circumstances such as dry seasons or winters (Denlinger, 2023; Halali et al., 2020; Nadeau et al., 2022). Tuning reproduction to cues that signal changes in environmental quality is key to fitness in seasonal habitats. This requires mechanisms that link sensitivity to relevant environmental signals and central cue processing with systemic signalling transducing this information and integrated or tissue-specific responses (van der Burg & Reed, 2021). Such seasonal plastic responses have been widely described across insects and especially in butterflies (Jones & Rienks, 1987; Kemp, 2001; Kopper et al., 2001; Lindestad et al., 2020; Radchuk et al., 2013), moths (Wadsworth & Dopman, 2015; Yamashita, 1996) and bees (Santos et al., 2018; Treanore et al., 2020).

Many tropical ecosystems exhibit strong seasonality in rainfall, which profoundly affects plant phenology and growth, and consequently places important constraints on insect reproduction (Feng et al., 2013). However, seasonal diapause is under-studied in the tropics compared to temperate areas (Denlinger, 2002, 2023; Halali et al., 2020; Nylin, 2013; Ragland et al., 2019; Saunders, 1933; Schebeck et al., 2024) (but see, Denlinger, 1986; Halali et al., 2021; Jones & Rienks, 1987; Kemp, 2001; Santos et al., 2018), in particular when considering tropical species’ proportion of global biodiversity. Recent work on diapause in the tropics has focused on the evolution of reproductive diapause associated with range expansion into African savannah regions (Halali et al., 2024; Tougeron, 2019). For instance, species across the Australian, Asian and African tropics display different forms of reproductive diapause: *Catopsilia pomona* (Jones, 1987) and *Eurema herla* (Jones & Rienks, 1987) show a preparatory and annually consistent cessation of reproduction, *Euploea core* (Daglish et al., 1986) and *Eurema laeta* (Jones & Rienks, 1987) have reproduction linked to rainfall, and members of the *Bicyclus* genus diapause in the dry season before or after mating (Halali et al., 2020). Outside of Lepidoptera there are also a range of descriptions of reproductive diapause in originally tropical species: *Drosophila melanogaster* has rapidly evolved a mild form of reproductive diapause in cosmopolitan North America with alleles that were already segregating in the ancestral tropical populations (Erickson et al., 2020) (and some argue that the species may have already had diapause in the African tropics (Zonato et al., 2017).

The growing concern is that climate change is making seasonal cues less reliable. The Neotropics are at a large risk of critical transitions in seasonality due to climate change (Nepstad et al., 2008), with models estimating that by 2050 there will be an increase in the number of consecutive dry days by 10-30 days across the Amazon forest system (Flores et al., 2024). Where species rely on consistent and reliable cues for timing of phenology, climate change-induced shifts in these seasonal patterns can lead to maladaptive phenotype-environment mismatches (Ashander et al., 2016; Stenseth et al., 2002; Thackeray et al., 2016; Visser & Both, 2005). This imposes strong selection on the extent and mechanistic architecture of plastic responses. Whether and how diapause will be able to evolve quickly enough to accommodate rapid anthropogenic climate change depends on many factors including alignment between axes of plasticity, fitness, and genetic variation (Chevin & Hoffmann, 2017; Noble et al., 2019; Riley et al., 2023). Understanding how plasticity has evolved in the past can enable predictions of how plasticity can evolve under climate change (Chirgwin et al., 2015).

While studies in temperate areas have revealed a great deal about mechanisms and evolution of diapause, it is unknown whether those insights also apply to reproductive plasticity in the tropics (Denlinger, 2023). Temperate diapause is tightly regulated by environmental cues (temperature and photoperiod), and it only ends when the climate becomes favourable (Denlinger, 2023; Kopper et al., 2001; Musolin & Saulich, 2012; Noer et al., 2022; Saunders, 1933). Whether it is a reproductive quiescence or diapause is also unknown: a reproductive *quiescence* is induced through a direct and immediate response to climatic changes, whereas a reproductive *diapause* is programmed to begin pre-emptively of annual seasonal fluctuations (Schebeck et al., 2024). However, these two phenomena are difficult to separate in natural populations even using molecular approaches (Lirakis et al., 2018; Poelchau et al., 2013). Evolutionary transitions between diapause and other forms of reproductive plasticity are also poorly understood; do they involve existing cues affecting more traits, or do traits gain sensitivity to new cues (Bhardwaj et al., 2020)? Work towards understanding these events in tropical areas is important because seasonality in tropical environments, and therefore regulation of insect behaviour, are shifting in the context of climate change.

Here, we analyse reproductive plasticity in Neotropical butterflies to reveal evolutionary patterns and climatic drivers of reproductive diapause in the tropics. We exploit a five-year butterfly monitoring time series from the southern Amazon, an important biodiversity hotspot and a major geographic origin of butterflies. We quantified reproductive status for 733 individual females of nine species across four subfamilies, focusing specifically on two closely related species from the genus Catonephele, and combined these with local weather station data. This enabled us to quantify how reproductive output varies across phylogenetic, seasonal, phenotypic and climatic axes, providing new insights into the evolution of insect diapause.

In the Amazon, there is a gradient in seasonality from the equator (non-seasonal) to higher latitudes (more seasonal). The south-west Amazon experiences around 3 months a year with less than 60mm of rain – a ‘tropical dry month’ as per the Köppen climate classification (Peel et al., 2007) Food and host plants are likely to become less abundant and lower quality during the dry season making it a less favourable environment (Braby & Jones, 1995). In many temperate regions, photoperiod, humidity, temperature, and host plant availability have all been shown to act as a partial cue to drive the onset of seasonal diapause (Goehring & Oberhauser, 2002; Klockmann & Fischer, 2019). The butterfly taxa studied here have diverged across the Amazon, representing around 79 million years of evolution (Chazot et al., 2021; Kumar et al., 2022). This provides an ideal model to study the evolution of reproductive plasticity in a seasonal tropical environment.

Our study aims to analyse reproductive plasticity in the tropics and how it varies along phylogenetic, seasonal, phenotypic and climatic axes. Specifically, we address the following questions: 1) How has reproductive diapause evolved in Nymphalid butterflies? 2) How does seasonal reproductive plasticity differ between two closely related *Catonephele* species? 3) What are the specific climatic drivers of seasonal reproductive plasticity in these species?

We observed a range of reproductive responses, even within genera, which suggests repeated changes in seasonal reproductive plasticity in multiple clades. Focusing on two closely related species of the subfamily Biblidinae, we observed *Catonephele acontius* to exhibit diapause during the dry season, while *C. numilia* does not. Despite their differences in reproductive plasticity, climate data analysis reveals that both species share maximum temperature as likely key driver of reproductive plasticity. Thus, evolution of diapause combines cue conservation with species-specific divergence of plastic traits, suggesting modular (step-by-step) evolution of reproductive plasticity. Importantly, our time series data enabled us to establish that the response to temperature is immediate, except for a small anticipatory role for photoperiod and minimum temperatures which may set up the temporal boundaries of direct temperature responses. Thus, diapause in our study species may share features both with direct stress responses and with anticipatory diapause as observed in other taxa, suggesting that evolutionary transitions between these different forms of plasticity may be gradual and modular.

## 2 | Material and methods

### 2.1 Study Area and Species’

Field collections of butterflies were conducted at Finca Las Piedras, a biological field station in southeastern Peru (Fig. s1) (lat. -12.226348°, lon. -69.112599°; ca. 250 masl) where we lead a long-term Lepidoptera monitoring and collection program. The lowland tropical rainforest site, characterised by a pronounced wet and dry season, offers an ideal setting for studying butterfly adaptations to a rapidly changing seasonal environment - specifically changes in reproductive strategy.

We first performed a broad analysis of reproductive dynamics in nine species across the Nymphalidae family, analysing species from the Biblidinae, Charxinae, Satyrinae and Nymphalinae subfamilies. We then focussed on reproductive dynamics in two butterflies *Catonephele acontius* and *C. numilia* (Lepidoptera: Nymphalidae: Biblidinae) with a 9-million-year divergence time (Hedges et al., 2015; Kumar et al., 2022). These species are sympatric across much of the Neotropics, including at the study site in Southeast Peru and are thought to share similar ecological niches, though very limited data is available. For instance, larvae of both species are known to feed on trees in the genus *Alchornea* (Euphorbiaceae) (Beccaloni et al. 2008, Woodson et al., 1967).

### 2.2 Specimen Collection

Standardised monthly trapping at FLP was carried out at ten trap stations in intact forest, each station with paired standard butterfly traps placed at ground level and at varying heights in the forest canopy and baited with fermented banana (Freitas et al., 2021). Trapping was conducted during five consecutive days within the first 10 days of each month, with the analyses presented in this paper covering June 2020 to January 2024. This protocol provides robust time-series data for the fruit-feeding species living in this seasonal environment.

To supplement the specimens collected as part of the long-term trapping study, additional collections were carried out at FLP in the dry season of 2023 (August – October). These included twenty banana-baited traps set up along a ∼2km stretch of forest trail, placed in light gaps near ground level. At the point of collection, geographic coordinates, date, time, sample ID and collector name were noted. Samples were then photographed, identified, and stored for future dissection in glassine envelopes.

### 2.3 Quantification of reproductive phenotypes

To determine reproductive status, we carried out dissections of adult female butterflies caught monthly throughout 2020-2024 (Fig. s2). We counted mature eggs (large/yolk-filled) and spermatophores in female butterflies and used these results to categorise reproductive activity based on the percent of diapausing individuals in the dry season, and whether the patterns were biased seasonally (see Table s1 in Supporting Information): no diapause (0-10%), seasonal diapause (25+%), limited nonseasonal diapause (10-25%) and finally a limited seasonal diapause (10-25%). The dissections were carried out using a Motic SMZ 160-TLED stereo microscope and standard dissection tools, following the procedure established by (Halali et al., 2020). The detailed protocol used in this study can be found in the supplementary information.

### 2.4 Climate data

Contemporary climate and weather data have been collected daily at Finca Las Piedras since July 2017 at chest height in primary forest (maximum/minimum daily temperature and daily precipitation). Long-term historic climatic data (1901-2021) was downloaded from the Climatic Research Unit Time Series (CRUTS) dataset 4.06 for the grid box -12.5, -69.5, - 12.0, -69.0 on the 24^th^ September 2024 (Osborn & Jones, 2014). This data was used for the Wavelet analyses and calculating Colwell’s Indices to test for climatic seasonality. Colwell’s indices give information about: environmental stability; repeatability of seasonal patterns; and the extent to which seasonality contributes to predictability (Colwell, 1974). We define the dry season as June to October, using a measurement of <60mm of total rainfall in the prior 30 days (Peel et al., 2007; Zeitschrift, 1884). This approach provides a stable estimation of the dry season across all four study years.

### 2.5 Statistical analyses

All statistical analyses and figure creation were carried out using R v4.2.2 (R Core Team, 2022), with the packages *WaveletComp v1.1* (Roesch A, 2018)*, hydrostats v0.2.9* (Bond N, 2022)*, ggplot2 v3.5.1* (Wickham H, 2016). For the wavelet analyses, the following functions within *WaveletComp* were used to calculate and then visualise the data: ‘analyze.wavelet’, ‘wt.image’ and ‘wt.avg’. The following parameters were used: loess.span = 0 (no smoothing), dt = 1 (evenly spaced monthly values), dj = 1/250 (fine scale resolution), lowerPeriod = 2 (months), upperPeriod = 32 (months), n.sim = 10 (simulations). The package *mondate v1.0* (“Mondate: Keep Track of Dates in Terms of Months,” 2010) was used to reformat the dates from the original database. *Hydrostats* was used to calculate Colwell’s indices (Table s2) using the ‘Colwells’ function. The following parameters were used: s = 11 (11 bins separates into 12 months), base.binning = 2 (logarithmic base binning), from = 0.5, by = 0.25, base.entropy=2 (binary logarithm).

To test differences in annual patterns of egg-laying, we used circular statistics with the packages *circular v0.5.1,* and *dplyr v1.1.4* (Lund & Agostinelli, 2024; Wickham H et al., 2025). Circular statistics relies on the conversion of cyclical data (e.g. months of the year) into angles. This approach returns a rho value which indicates how clustered the data are (0 = nearly uniform across the year, 1 = highly concentrated in a few months) and a mean angle, around which the data are clustered. Due to unknown and likely uneven sampling effort, we took 1,000 bootstrap resamples of butterflies within each month, which gave us a 95% confidence interval of rho for the egg-laying of each species.

To test the correlation between our phenotypic and climatic variables, we used both linear regressions and Spearman’s rank correlation. These correlational analyses used weather data collected and recorded daily at Finca Las Piedras which was then averaged per month. All raw data and R scripts can be found in the Git repository linked in the supporting data.

## 3 | Results

Given the range of existing definitions of reproductive diapause, in this paper we compare reproductive activity for all species through multiple measurements, specifically egg number within reproductive females (defined as those holding at least one mature egg), the proportion of individuals without eggs (diapausing) and the proportion of individuals without spermatophores (non-mating). Some butterflies could lack eggs due to other factors (e.g. health/age) which we are unable to quantify.

### 3.1 Pronounced dry-wet seasonality of study area

Although there are pronounced differences in the climate between the wet and dry season in this area of the Amazon, there is no well-defined boundary. To demonstrate that there is a periodic (i.e. annual) change in climatic variables, we performed a wavelet analysis of two main climatic variables (mean temperature and precipitation) using historical data from 1921-2020 (Fig. s3). Wavelet analyses are used for quantifying or identifying periodicity of fluctuations in environmental or other variables and here showed significant and coordinated annual periodicity for both temperature and precipitation. Furthermore, Colwell’s indices (Colwell, 1974) (Table s2) suggests that there is medium to high environmental stability at FLP, and that seasonality contributes to predictability, meaning that it is appropriate to define a wet and dry season. This analysis justifies the use of the Köppen climate classification: <60mm in the previous 30 days to define a dry season day. It is important to note, however, that these analyses are based on modelled historical data (CRUTS), covering an area of approximately 5km^2^.

### 3.2 Variation in reproductive plasticity across the Nymphalidae family

We show that changes in reproductive status during the dry season are common across Nymphalids (Fig. 1). Specifically, *Bia rebeli* of the Satyrinae subfamily does not show any diapause, whilst in Charaxinae there are different species demonstrating either a seasonal or non-seasonal diapause (acknowledging smaller sample sizes). Both *Colobura* species in the Nymphalinae subfamily demonstrate a full seasonal diapause, but in the Biblidinae subfamily it is only *Catonephele acontius* that shows a seasonal diapause, whilst *Nessaea obrinus* show a limited nonseasonal diapause and *Catonephele numilia* does not show any seasonal diapause.

**Figure 1.**
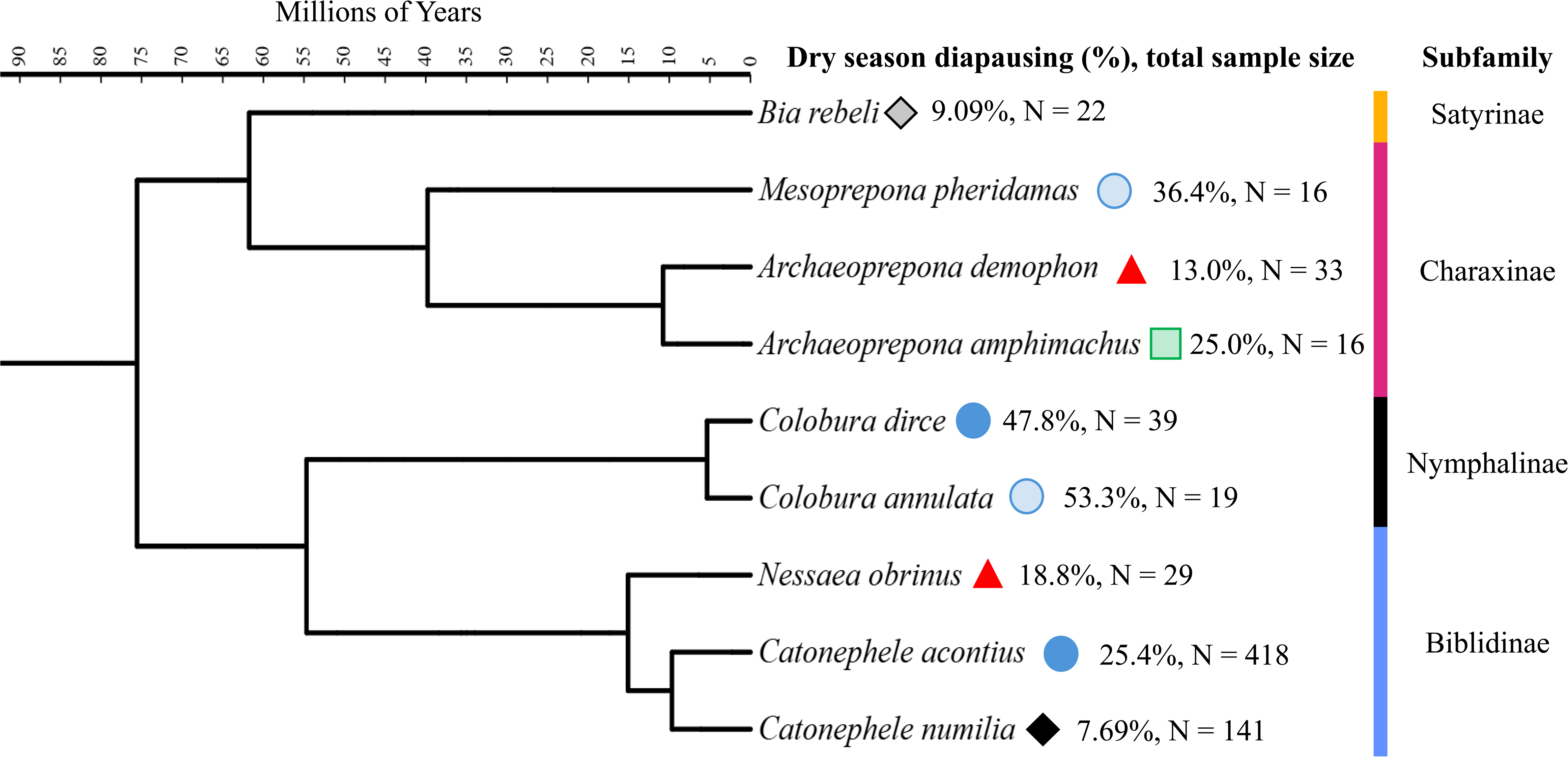
Variation in seasonal reproductive plasticity across 79 million years of Nymphalid butterfly evolution in a neotropical rainforest. We phenotyped reproductive status by counting eggs for 733 individual females of 9 species (in 4 subfamilies) collected in dry and wet seasons over five years. Females without any eggs were categorised as diapausing, and numbers were compared between dry and wet seasons to classify species into 4 categories of seasonal reproductive plasticity: no diapause (black diamonds), seasonal diapause (blue circles), limited diapause but nonseasonal (red triangles) and finally limited seasonal diapause (green squares). The proportion of females diapausing during the dry season, and the total sample size for that species are noted. Lighter fill colour shades indicate a small sample size (<20). Based on phylogeny from (Chazot et al., 2021; Letunic & Bork, 2024). A more detailed breakdown of egg number patterns can be found in supplementary table 1.

### 3.3 Seasonal diapause in *C. acontius* but not *C. numilia*

The two *Catonephele* species showed markedly different seasonal responses in egg production (Fig. 2). The egg number distribution for *Catonephele acontius* differs significantly between the wet and the dry season, with a significantly higher proportion of diapausing individuals during the dry season (25.4% of 177 individuals having no eggs) compared to the wet season (8.7% of 241 individuals; X-squared = 20.192, df = 1, p-value = 7.003e-06). In contrast, while a small proportion of C. numilia individuals were also diapausing, there was no difference between dry and wet seasons (wet: 9.21%, n =76; dry: 7.69%, n =65; X-squared = 0.00037338, df = 1, p = 0.9846).

**Figure 2.**
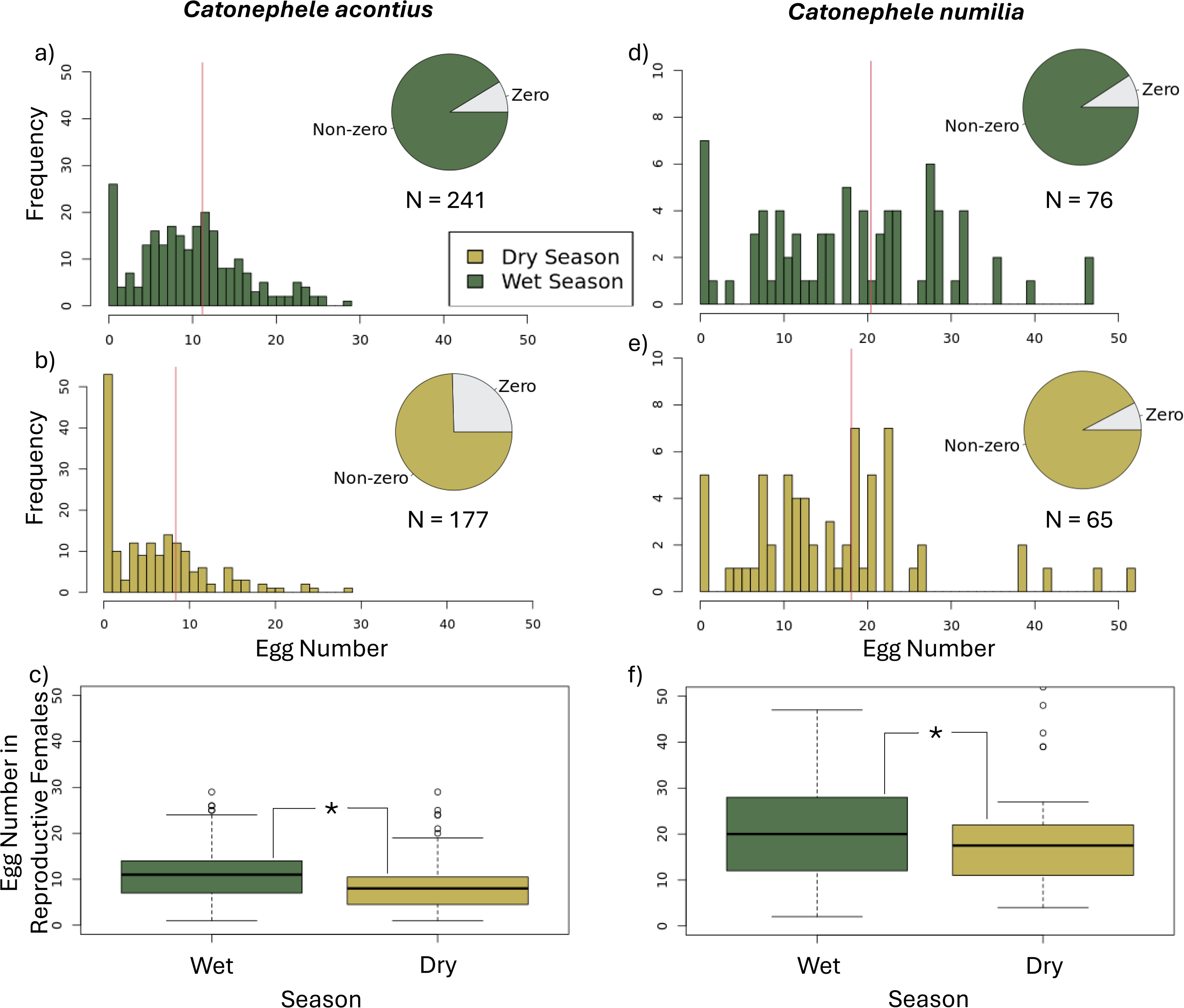
**Dry season induces a marked shift in diapause strategy in *Catonephele acontius* but not in *C. numilia.*** Distribution of egg number in a,b,) *C. acontius* and d,e) *C. numilia* in wet and dry seasons respectively. Histograms show a zero-inflated, positively skewed normal distribution of egg number in both, but the number of diapausing females strongly increases between wet and dry seasons in *C. acontius* and not *C. numilia.* The average number of eggs (red vertical lines in a, b, d, e) for reproductive females reduces significantly from the wet season to the dry season in c) *C. acontius* (*Wilcoxon rank sum test one-sided, W = 10063, p-value = 6.818e-07) and f) *C. numilia* (*Wilcoxon rank sum test one-sided, W = 1706, p-value = 0.04287). A comparison with a randomly selected equal sample size demonstrates similar overall patterns (Fig s4).

However, when considering only reproductively active females (i.e. those with at least one egg), we observed a significant reduction in egg number between wet and dry seasons for both *C. acontius* (Wilcoxon rank sum test one-sided, W = 10063, p-value = 6.818e-07) and *C. numilia* (Wilcoxon rank sum test one-sided, W = 1706, p-value = 0.04287) (Fig. 2). This suggests a common strategy for reproductive females to lower reproductive output during the dry season, despite the clear difference in the strategy of diapausing individuals.

We observed a less pronounced seasonal change in reproductive activity (spermatophore number) in comparison to reproductive output (egg number). Specifically, we observed how most individuals of both species hold 1-3 spermatophore, and a small proportion of the population being unmated (Fig. 3). *C. acontius* caught in the dry season were more likely to be unmated (not carrying spermatophores) than those caught in the wet season (wet: 4.98%, n =241; dry: 12.27%, n =177, X-squared = 4.9068, df = 1, p-value = 0.02675). However, the same pattern was not found in *C. numilia* (wet: 4.62%, n =76; dry: 2.63%, n =65, X-squared = 0.031743, df = 1, p-value = 0.8586) (Fig. s5). A reduction of 13.32% (wet: 1.65, dry: 1.43) was seen in average spermatophore number for *C. acontius* in the dry season compared to the wet season (Wilcoxon rank sum test one-sided, W = 16968, p-value = 0.005701). No such difference was observed in *C. numilia* (Wilcoxon rank sum test one-sided, W = 2917, p = 0.9729).

**Figure 3.**
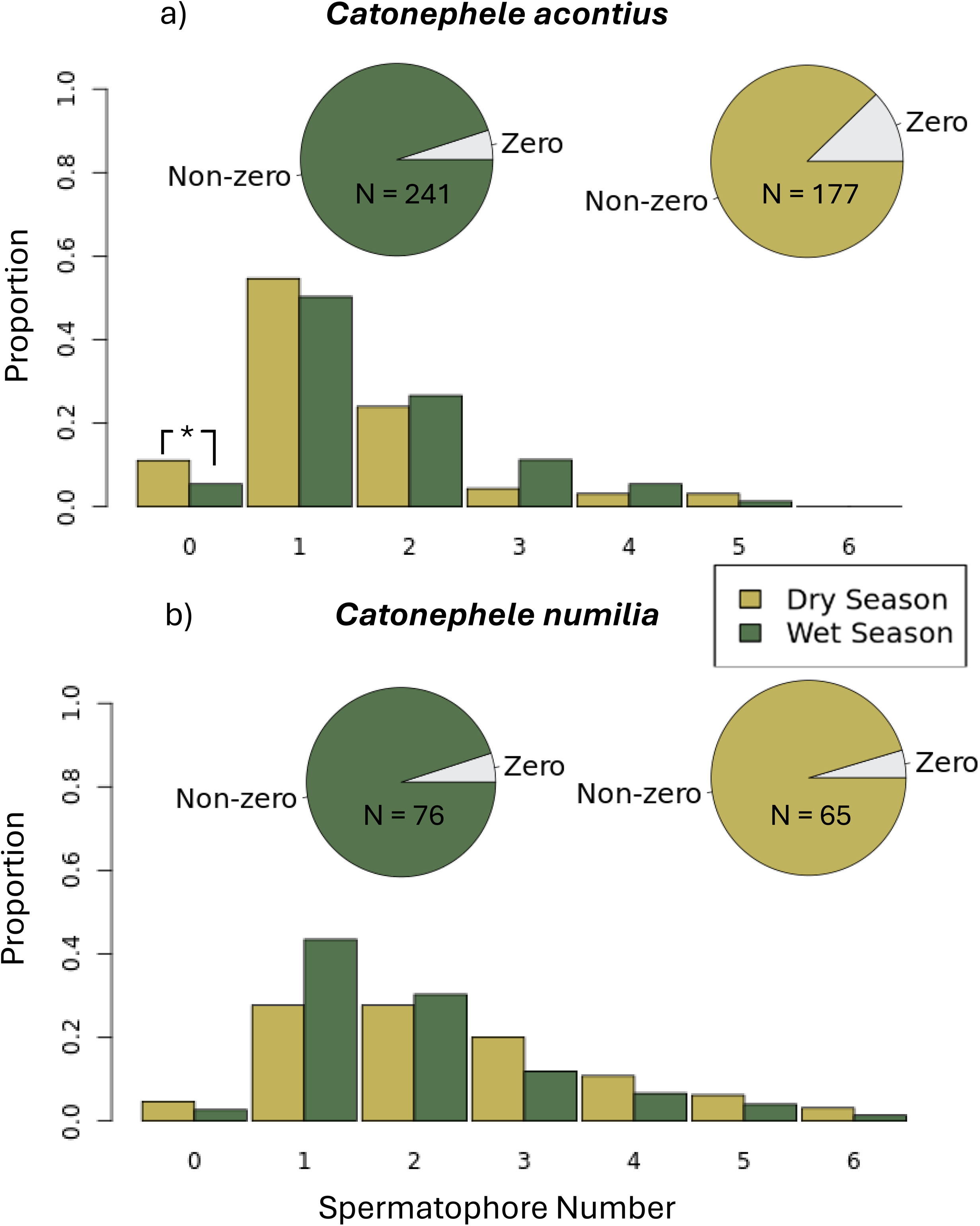
**Dry season induces muted change in reproductive activity in *C. acontius* but not in *C. numilia.*** Distribution of spermatophore number in a) *C. acontius* and b) *C. numilia* in wet and dry seasons respectively. Pie charts show that the proportion of mated individuals (with spermatophores) varies little between the seasons and between the species, with most individuals holding one to three spermatophore at any time. There is a minor, yet significant, increase in the proportion of non-mated (zero spermatophore) individuals in *C. acontius* in the dry season (*wet: 4.98%, n =241; dry: 12.27%, n =177, X-squared = 4.9068, df = 1, p-value = 0.02675).

Monthly time-series data for egg number throughout the year allows us to examine intra-annual fluctuations in reproductive output in more detail (Fig. 4). Both species demonstrate a non-uniform distribution (Rayleigh test, *C. acontius:* z = 0.22, p = 1.1^-75^, *C.* numilia: z = 0.109, p = 1.4^-13^). However, *C. acontius* has a more distinct seasonal response in egg production with egg-laying more clustered in the wet season for *C. acontius* (median rho = 0.22, 95% CI: 0.178–0.261, mean month = 11.2 (November), bootstrapped n = 1000), than in *C. numilia* (median rho = 0.109, 95% CI: 0.056–0.162, mean month = 9.93 (September), bootstrapped n = 1000).

**Figure 4.**
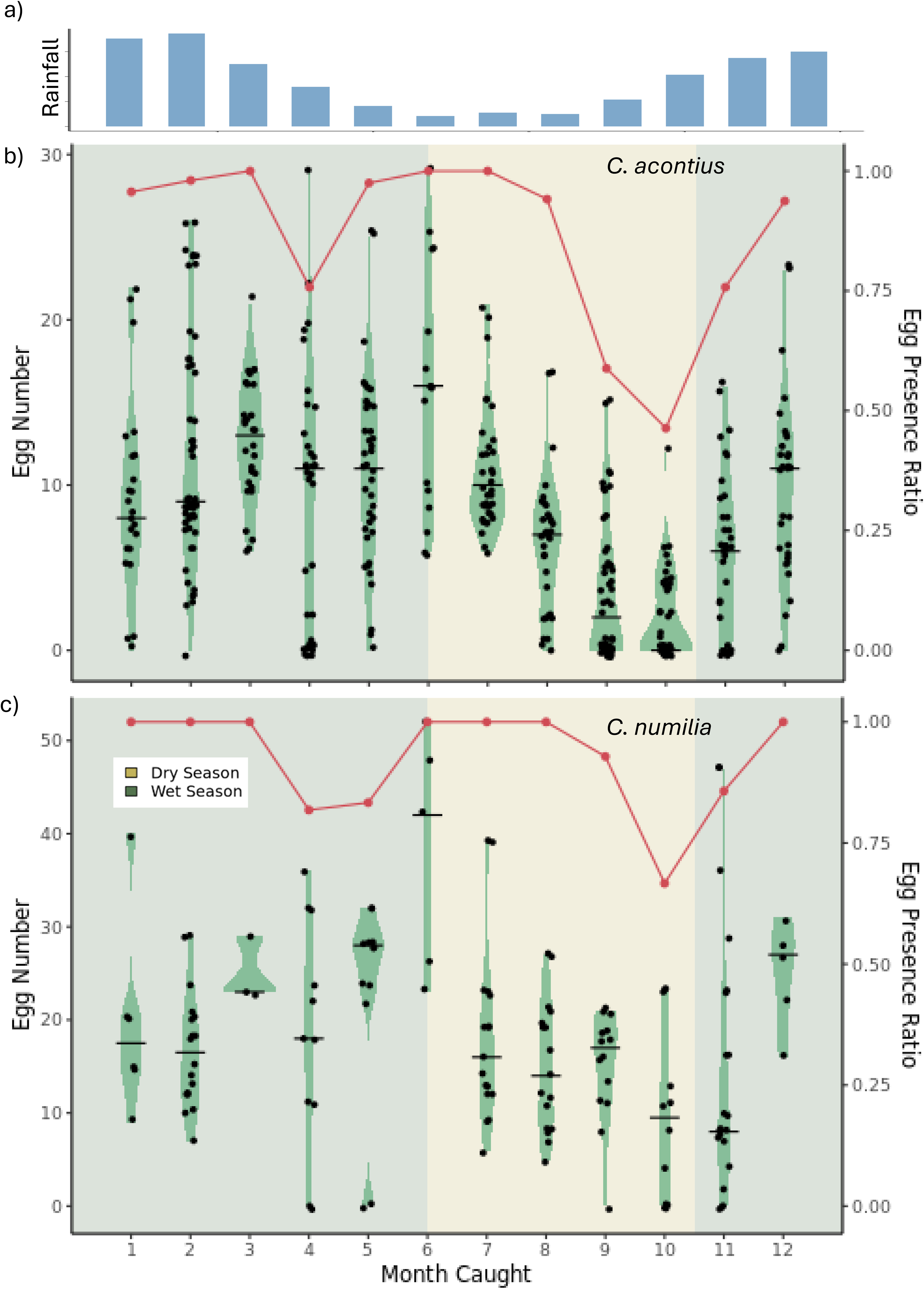
A decrease in average egg number during the dry season in both species, but a seasonal increase in diapausing individuals occurs only in *C. acontius,* and not in *C. numilia.* The seasonality in rainfall is indicated by a) average daily precipitation per month (January to December) and the seasonal background colours in b and c. Violin plots are created with data from the dissections of b) *C. acontius* and c) *C. numilia* in FLP aggregated over 2020-2024 with the black horizontal line representing the median value for that month. Thickness of the violin represents density of data. The sample size for each month can be found in Table s3. The red line tracks the proportion of reproductive individuals (holding at least one egg), demonstrating the change in diapausing strategists throughout the year. There is low interannual variability in these time series data (Fig. s6) suggesting that the behaviour is consistent, at least during this study period.

### 3.4 Climate-phenotype relationships

To understand how the environment drives seasonal reproductive dynamics of Catonephele acontius, we examined the correlation of each phenotypic variable (proportion of diapausing females; average egg number of reproductive females; proportion of unmated females) with environmental variables (rainfall, temperature, photoperiod), using monthly averages for each variable (Fig. 5). There is a strong relationship of maximum temperature with all three phenotypic variables compelling an immediate response during the dry season (egg number: R^2^ = -0.697, S = 506, p-value = 0.005253; diapause proportion: R^2^ = 0.696, S = 46.901, p-value = 0.0007043; unmated proportion: R^2^ = 0.468, S = 108.76, p-value = 0.0316) (Table s4).

**Figure 5.**
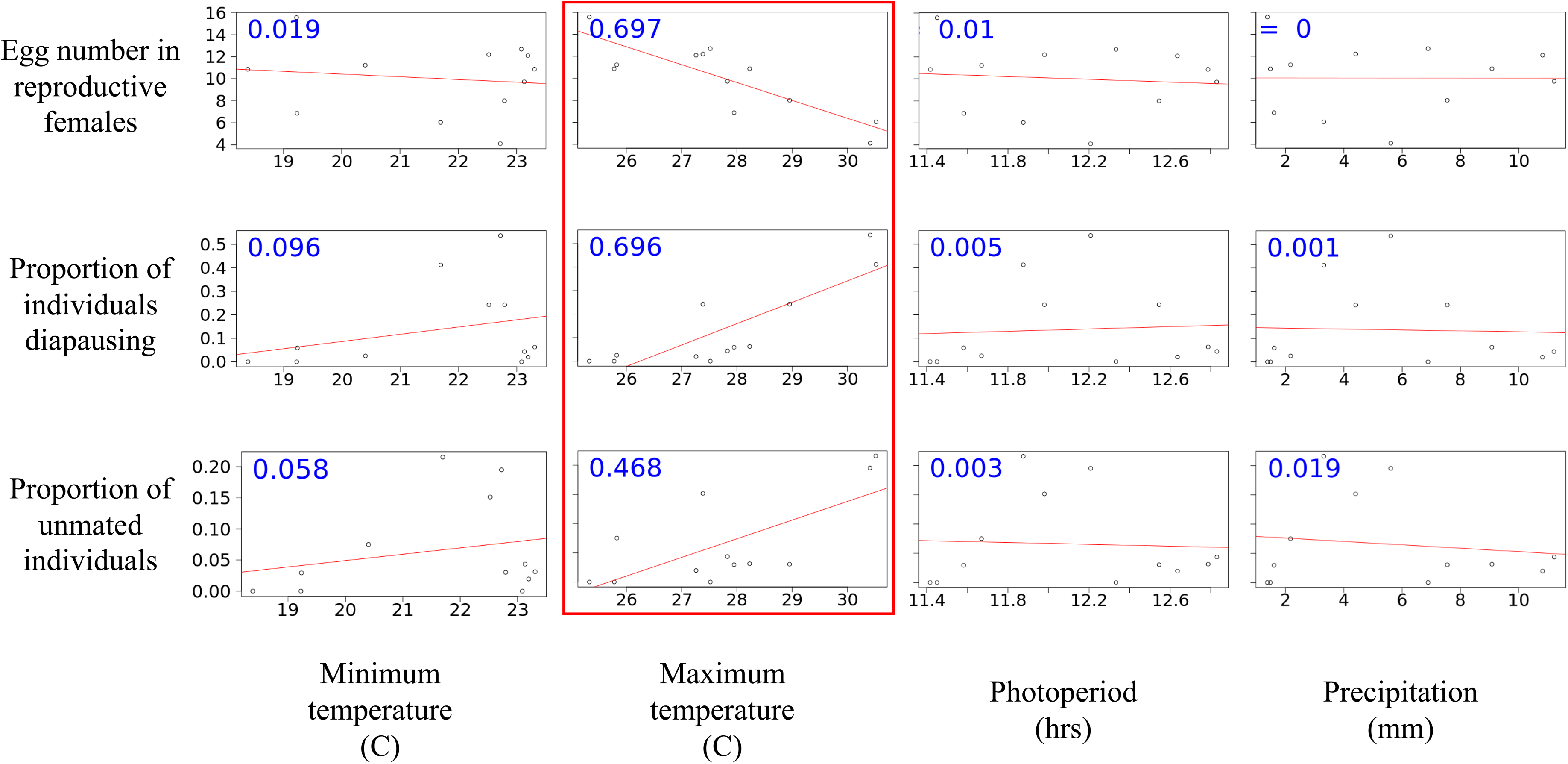
Maximum temperature correlates strongly with all phenotypic variables. Results from a correlational analysis of monthly values between four climate variables (maximum temperature, minimum temperature, photoperiod and precipitation) and the three phenotypic variables (egg number in reproductive females, proportion of individuals diapausing, and proportion of unmated individuals). Blue numbers are r^2^ values (Spearman’s rank correlation coefficient), whilst red lines are linear model regressions.

In the same analysis for *C. numilia,* maximum temperature was also the best correlate for the average egg number in reproductive females, suggesting a potentially shared underlying mechanism (R^2^ = -0.435, S = 468, p-value = 0.0301).

This also tested time-shifted correlations between phenotypes and climate 1 or 2 months earlier (testing if and how phenotypic changes precede seasonal shifts in a potential anticipatory response) or 1 or 2 months later (testing if and how potential environmental cues precede phenotypic changes) (Fig. 6). The two-month time lag is based on the natural lifespan of the two species (Gonzalez, 2024). Across photoperiod, minimum temperature, and daily rainfall, there are significant correlations with egg number in reproductive females 2 months earlier (minimum temperature: R^2^ = -0.461, S = 494, p-value = 0.01; photoperiod: R^2^ = -0.574, S = 500, p-value = 0.007353; precipitation: R^2^ = -0.439, S = 492, p-value = 0.01102) and 2 months later (minimum temperature: R^2^ = 0.638, S = 68, p-value = 0.005897; photoperiod: R^2^ = 0.427, S = 68, p-value = 0.005897; precipitation: R^2^ = 0.414, S = 68, p-value = 0.005897). This suggests a complex relationship between the climate and the onset and termination of these dry season phenotypes, with maximum temperature a key variable.

**Figure 6.**
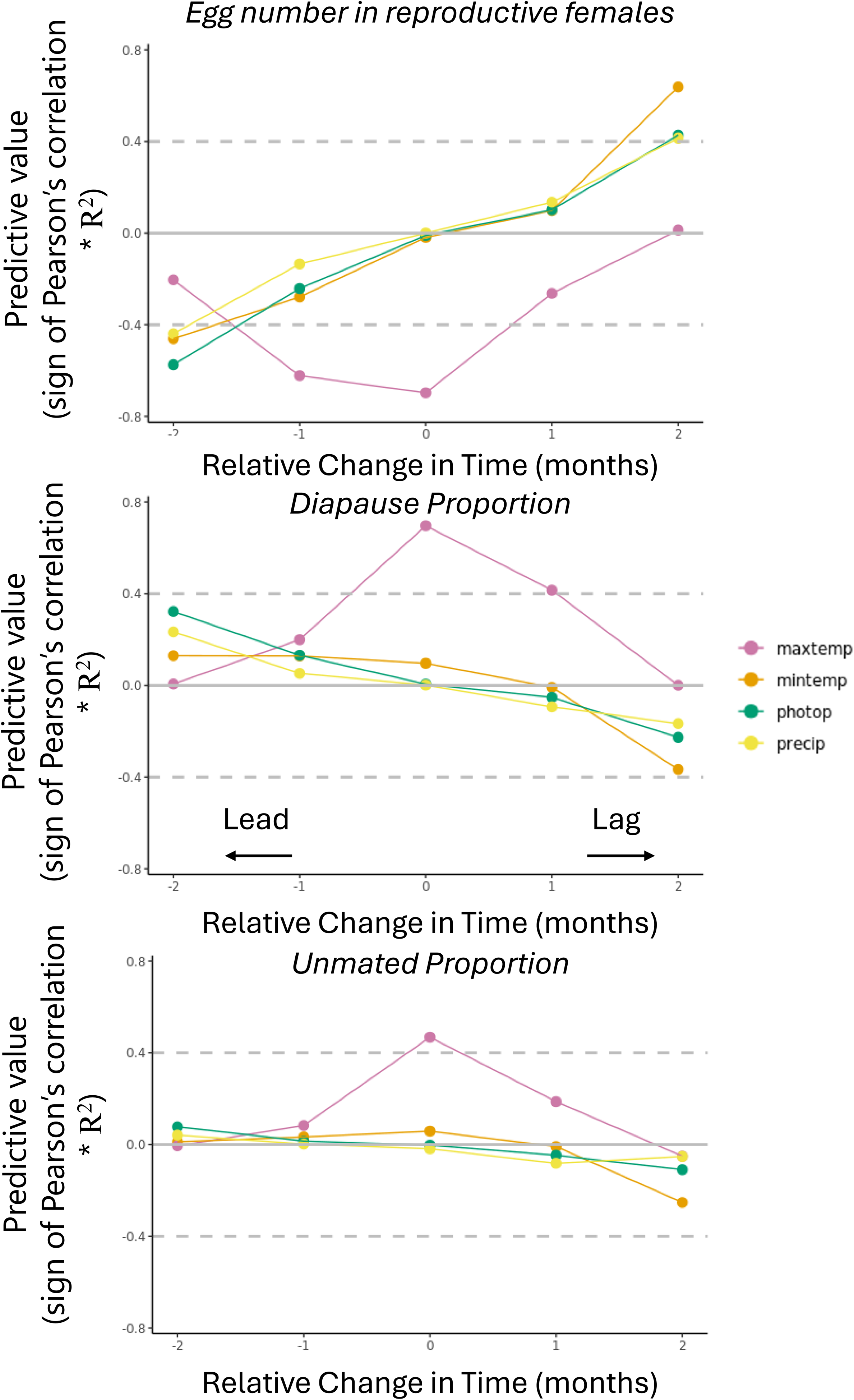
Maximum temperature has a unique relationship with phenotypic variables when considering different sensing and response mechanisms in *C. acontius*. These plots demonstrate the predictive value of environmental variables for reproductive traits (sign of Pearson’s correlation * R^2^) with different relative changes in time (+/- 2 months). Minimum temperature, photoperiod and precipitation all follow similar patterns, having the most predictive value at plus or minus two months. Maximum temperature, on the other hand, is unique in having the strongest predictive value with no temporal change, in other words having an immediate effect on the phenotype in question.

In summary, we observe that maximum temperature has a completely different relationship with these life history variables, compared to minimum temperature, photoperiod, and rainfall. The other climatic variables show associations with the egg number in reproductive females in both the two-month lead and two-month lag scenarios, whilst maximum temperature is only strongly predictive when considered to be moving in tandem with all three reproductive phenotypes (Fig. 6).

## 4 | Discussion

Here, we leverage a rare long-term butterfly monitoring dataset from the Neotropics, an important biodiversity hotspot and a major geographic origin of butterflies (Kawahara et al., 2023). We use detailed abdomen dissections for 733 individual females of 9 species representing around 79 million years of evolution and combine these with local weather station data. This enabled us to quantify how reproductive output varies across phylogenetic, seasonal, climatic and phenotypic axes, providing new insights into the evolution of insect diapause, and its potential role in adaptation to climate change. Our work suggests that the modularity of reproductive plasticity may facilitate a stepwise evolution of diapause, which will alter populations’ ability to persist in the face of climate change. Our climate analysis adds an important new angle to help us to predict how species may respond to a changing climate. Using long-term field data, we fill a key gap in our understanding of seasonal behaviour in Neotropical insects.

Sampling 4 subfamilies across the Nymphalid phylogeny (divergence times spanning 10-79 MYA), we observe phylogenetically widespread occurrence of diapause, including seasonal and nonseasonal diapause. This suggests repeated changes in reproductive plasticity in multiple clades and would make sense given a tropical origin of reproductive diapause. This finding is important for two key reasons: firstly, because the Neotropics are likely to experience intense changes under future climate change (Flores et al., 2024). Secondly, these results add the Neotropics to the current geographical range where reproductive diapause has been found, a novel but expected finding given the instances of this trait across other tropical regions (Braby & Jones, 1995; Denlinger, 1986, 2023; Halali et al., 2021, 2024; Jones & Rienks, 1987; Kemp, 2001; Tauber & Tauber, 1981; Tougeron, 2019).

Next, using two *Catonephele* species as a model, we show that one species, *C. acontius*, exhibits diapause during the dry season, while *C. numilia* does not. In *C. acontius* the proportion of diapausing and unmated females significantly increases during the dry season while we observe no such response in *C. numilia*. This indicates divergent evolution in seasonal reproductive strategy in this genus that has happened since their last common ancestor 9MYA (Hedges et al., 2015; Kumar et al., 2022). At the same time, we observe seasonal shifts in other aspects of reproductive activity that are shared between the species, suggesting a conserved component of their mating strategy. Specifically, reproductive (non-diapausing) females of both *Catonephele* species show a modest reduction in egg production during the dry season. Importantly, both the conserved and species-specific seasonal shifts in reproductive plasticity are strongly correlated with predictable seasonal increases in maximum temperature. This suggests that existing environmental cues can drive new phenotypic responses, via shared sensing and regulatory systems. Under climate change, such a direct response to temperature could lead to rapid induced shifts in mating strategy, but these may be maladaptive due to phenological mismatches unless climatic correlations stay constant (Walker et al., 2019). Together, our results provide evidence for dramatic seasonal shifts in a suite of reproductive traits in Neotropical rainforest insects, with species-specific and conserved components driven by shared environmental cues.

Species of tropical savannah-dwelling butterflies have been found to undergo a seasonal reproductive diapause in both Africa and Australia (Braby & Jones, 1995; Halali et al., 2020; Jones & Rienks, 1987; Kemp, 2001); we are highlighting one of the first examples of seasonal diapause in tropical forest species. An adaptation like reproductive diapause may consequently allow these butterfly species to expand their ranges into the Cerrado (Neotropical savannah), like in *Bicyclus* butterflies (Brakefield, 2010). This niche expansion is an important evolutionary process, especially in the context of climate change, given that Neotropical dry seasons are predicted to become longer and more severe, leading to expansion of the Cerrado across the continent if tipping points are reached (Flores et al., 2024). As rainforest becomes more savannah-like, species with pre-existing diapause responses may benefit. However, this critically depends on the reliability of seasonal cues – which is predicted to decrease with climate change (Nepstad et al., 2008).

Our climate data analysis revealed that maximum temperature is the key driver of reproductive plasticity for both our study species. In well-studied temperate instances of diapause, it is common to see temperature and photoperiod playing a role in the timing of developmental arrests (Goehring & Oberhauser, 2002; Klockmann & Fischer, 2019). Our results suggest that this Neotropical seasonal diapause – across a suite of correlated traits – is induced by a single and direct response to maximum temperature, with possible ‘setting up’ via photoperiod. Temperature has been linked to seasonal plasticity when coupled with changes in photoperiod (Mallick et al., 2024; Schebeck et al., 2024; Sgrò et al., 2016) and has been well studied in temperate Lepidoptera (Lindestad et al., 2020; von Schmalensee et al., 2024).

Interestingly, our time series analysis of long-term climatic data (Fig. 5) indicates that the reproductive shift is a direct, immediate response to elevated temperature, rather than an anticipatory response to the upcoming dry season. This is like other types of stress responses (e.g. heat stress in Denlinger, 2002; Gong et al., 2013; Yocum et al., 1998) where a reduction in environmental quality directly arrests reproduction. The direct response to temperature suggests that diapause exists along a spectrum of reproductive plasticity that includes immediate stress responses. Together with our observation of repeated evolution of reproductive plasticity across a broad phylogenetic distribution (Fig 1), this may indicate that evolutionary transitions between these different forms of plasticity are gradual and modular.

What is unknown is whether maximum temperature is mainly acting as the selective agent (sensu Nijhout, 2003). Another (non-exclusive) role of temperature may be that it acts as a predictive cue to induce diapause, which could have evolved as an adaptation to other harsh aspects of the dry season – for example a reduction in host plant quality or extremely low rainfall. Photoperiod is one of the key diapause inducing environmental cues used by many temperate insects (Jin et al., 2024; Pruisscher et al., 2021; Sgrò et al., 2016). Within complex climatic conditions, the correlations between climatic variables become important for producing an appropriate phenotype matched to the selective environment. As climate change may rapidly alter the relationships between climatic variables, there is a concern that phenotype-environment mismatches will occur (Walker et al., 2019). However, in the absence of manipulative experiments it is difficult to disentangle environmental variables as part of either the inductive or selective environment.

The responses of a suite of correlated traits to correlated changes in the environment in both *C. acontius* and *C. numilia* could suggest shared developmental, physiological and genetic pathways, as well as a shared evolutionary history. Diapause likely evolved by integrating new output to existing modules of cue induction and phenotype regulation. This is supported by other models of reproductive diapause in temperate insects, where similar groups of hormones and signalling systems are implicated in creating a diapause response that involves multiple coordinated traits. The ‘toolbox’ of genes underlying the diapause response consists of photoperiod sensing (Green & Kronforst, 2019; Musolin & Saulich, 2012; Saunders, 1933; Sgrò et al., 2016), insulin signalling (Green & Kronforst, 2019; Nylin, 2013; Sim & Denlinger, 2013; Tatar & Yin, 2001) and select hormones, with Juvenile Hormone, PTTH and ecdysone being the most important (Denlinger, 2002; Marden et al., 2008; Saunders, 1933; Telfer, 2009). Suites of correlated traits tend to be regulated by the same hormones (Van Bergen et al., 2017) and respond to the same (often multivariate) environmental cues (Singh et al., 2019). This impacts the evolutionary pathways of these modules, with selection on phenotypes favouring modular genetic systems – including hormonal regulation (Cox, 2020; Cox et al., 2016; Murren, 2012). This could therefore be an explanation for the evolution of diapause in these Neotropical insects: the inductive role of maximum temperature may have been co-opted from an existing stress response that downregulates egg production and mating behaviour under poor conditions. However, it is difficult to know if this true diapause trait is, as suggested, a discrete phenotype or if it is a continuous reaction norm hitting the minima of egg production (Nijhout, 2003; Plaistow & Collin, 2014; Schlichting, 1989).

Notably, not all *C. acontius* females go into complete reproductive arrest in the dry season. At the peak of the dry season, 50% of females continue to reproduce, albeit it at reduced rate (Fig. 4). The fitness benefits of completely repressing reproduction may outweigh the costs only under some conditions or only for some individuals. This may represent a bet hedging strategy that reduces variance in fitness in variable environments (Overton & Sharkey, 2021). Alternatively, if the ability to undergo diapause is genetically determined as it is in other insects (e.g. Erickson et al., 2020), this genetic variation could be maintained through balancing selection driven by inter-year variation in environmental quality.

Given the distinct reproductive dynamics of these two species, it is relevant to consider the ecological similarities and differences between them. Both species share similar ranges across South America, with *C. numilia* having a slightly more northerly range into Central America (*INaturalist*, 2024). They share the same group of host plants within the *Alchornea* genus and are likely to both specialise on *A. latifolia* in the south-eastern ranges of Peru (Alberto Muyshondt, 1975). They have similar developmental timings and adult lifespans in captivity (Gonzalez, 2024). One difference that we have found is their abundance patterns in Finca Las Piedras (Fig. s7). *C. acontius*, the diapausing species, has a higher abundance throughout the year, and has two major peaks – one in April at the peak of the wet season, and one in August, at the height of the dry season. *C. numilia* on the other hand, shows a much more constant abundance throughout the year, with just one peak during the dry season. This is consistent with different life-history strategies between the diapausing and non-diapausing species.

Reproductive output is something that changes throughout a butterfly’s life-history, and it therefore may be that the age of the individuals sampled acts as a confounding variable in recognising trends in reproductive output. Without mark and recapture, wild-caught butterflies are difficult to age but progress has been made on methods of using wing damage as a proxy (Le Roy et al., 2019; Lehnert, 2010; Molleman et al., 2020). Future work formalising this and the possible impact on our analyses would be beneficial. Similarly, due to these individuals being wild caught, we cannot experimentally control environmental effects, and we therefore base our inductive cue conclusions on correlational analyses of climate data. To untangle various correlated climatic variables, future work should focus on long-term collections of the same species in a Neotropical environment that is seasonal in temperature and photoperiod, but not in rainfall and host-plant quality.

Our work reveals that reproductive plasticity is phylogenetically widespread in the Neotropics. Our analyses also suggest that diapause in our study species may share features both with direct stress responses and with anticipatory diapause as observed in other taxa, suggesting that evolutionary transitions between these different forms of plasticity may be gradual and modular. The exact evolutionary outcomes likely depend on the interplay between large-scale gradients in seasonality, and local inter-year variability in environmental quality. Ongoing work on ecological, evolutionary and genetic drivers of diapause in tropical species promise to further clarify long-standing questions in the evolution of diapause in complex tropical environments.

## Availability of supporting data

All raw data and metadata supporting the manuscript and R code for all analyses are available on figshare: https://doi.org/10.6084/m9.figshare.30227872.v1 (Hicks et al., 2025).

## Acknowledgements

MH is funded by a QMUL PhD scholarship. He received additional funding from the Genetics Society, QMUL and NERC Environmental Omics Facility (NEOF) (grant NEOF1704).

VO is funded by QMUL and a UKRI Future Leaders Fellowship (MR/V024744/2).

GRG acknowledges the financial support of Wild Green Future. The authors thank Peru’s Servicio Nacional Forestal y de Fauna Silvestre (SERFOR) for permission to conduct field and laboratory research (permit no. D000443-2021-MIDAGRI-SERFOR-DGGSPFFS).

We also thank Alexander, Savi and other members of the Alliance for a Sustainable Amazon for their help in collections, and particularly the Lepidoptera team.

There are no conflicts of interest.

## Author Contributions

MH, GG, and VO conceptualized the study. JK carried out the initial pilot study under the supervision of GG. MH, LR and ZE carried out the fieldwork, supervised by GG and VO. Data analysis was done by MH, supervised by both SH and VO, and writing of the manuscript was done by MH with input and supervision from VO. All authors contributed to the revision of the final manuscript.

The data and scripts supporting the results will be archived in an appropriate public repository and the data DOI is included at the end of the article.

